# Mycobacterial EtfD contains an unusual linear [3Fe-4S] cluster and enables β-oxidation to drive proton pumping by the electron transport chain

**DOI:** 10.1101/2025.05.14.654035

**Authors:** Gautier M. Courbon, Vadim Makarov, Stewart T. Cole, Dirk Schnapinger, Sabine Ehrt, John L. Rubinstein

## Abstract

In mycobacteria, the protein EtfD is thought to link β-oxidation of fatty acids with the electron transport chain, two processes that have attracted attention as targets for therapeutics to treat tuberculosis (TB) and other mycobacterial infections. It has been proposed that targeting β- oxidation could shorten treatment duration by killing non-replicating *Mycobacterium tuberculosis* within granulomas in the lungs. Here we show that *Mycobacterium smegmatis*, a fast growing and nonpathogenic model for energy metabolism in *M. tuberculosis*, relies on EtfD for extracting energy from β-oxidation. Electron cryomicroscopy allowed structure determination of *M. smegmatis* EtfD, revealing an unusual linear [3Fe-4S] cluster that has not been seen in other protein structures, but which resembles the catalytic noncubane [4Fe-4S] clusters in heterodisulfide reductases. The structure suggests how EtfD transfers electrons from β-oxidation to the electron transport chain. We devised an assay that couples EtfD activity to a fluorescent readout of proton pumping by the electron transport chain, which can be used to identify compounds that block mycobacteria from using β-oxidation to power oxidative phosphorylation.

## Introduction

During infection, *Mycobacterium tuberculosis*, which causes the disease tuberculosis (TB), can survive within structures in the lung formed by immune cells and known as granulomas (Russell, 2001; Barry et al., 2009). In the lipid-rich and hypoxic environment of the granuloma, the bacterium undergoes metabolic remodeling that allows it to enter a persistent non-replicating state associated with tolerance to drugs, necessitating long treatment durations for infections (Gomez and McKinney, 2004; Boshoff and Barry, 2005; Ehrt et al., 2018). The discovery and clinical success of the adenosine triphosphate (ATP) synthase inhibitor bedaquiline, which kills both replicating and non-replicating *M. tuberculosis in vitro* (Koul et al., 2008), has validated mycobacterial energy metabolism as a target for the development of new drugs with the potential to shorten TB treatment (Andries et al., 2005; Deshkar et al., 2022). The use of cholesterol and fatty acids as a carbon source in granulomas, where other nutrients are scarce, could play an important role in persistence (Boshoff and Barry, 2005; Russell et al., 2009). Therefore, disrupting *M. tuberculosis* lipid catabolism has been proposed as a strategy to target persistent mycobacteria, which could also reduce treatment duration (Wilburn et al., 2018; Beites et al., 2021).

Metabolism of fatty acids to release energy uses the β-oxidation pathway. This pathway is tightly integrated with the central carbon metabolism of the Krebs cycle (**Fig. 1A**). In β- oxidation, fatty acids are activated by an acyl-CoA synthetase that attaches coenzyme A (CoA) to the first carbon of the acid, forming a fatty acyl-CoA. The fatty acyl-CoA is subjected to multiple iterations of processing by acyl-CoA dehydrogenases, enoyl-CoA dehydratases, β-hydroxyacyl-CoA dehydrogenases, and thiolases. Each iteration shortens the fatty acid’s aliphatic chain by two carbons, releases a molecule of acetyl-CoA, reduces a soluble nicotinamide-adenine dinucleotide molecule (NAD^+^) with two electrons to form NADH, and reduces a protein-bound flavin adenine dinucleotide (FAD) cofactor with two electrons to form FADH2. The NADH and FADH2 are oxidized by the electron transport chain (ETC), which ultimately establishes an electrochemical proton motive force (pmf) across the plasma membrane. The pmf is used by ATP synthase to generate ATP. The acetyl-CoA from β-oxidation enters the Krebs cycle, leading to the reduction of either three NAD^+^ molecules and one protein-bound FAD, or two NAD^+^ molecules and two protein-bound FADs, depending on which malate oxidizing enzyme is used by the organism (Houten and Wanders, 2010; Harold et al., 2022).

**Figure 1.**
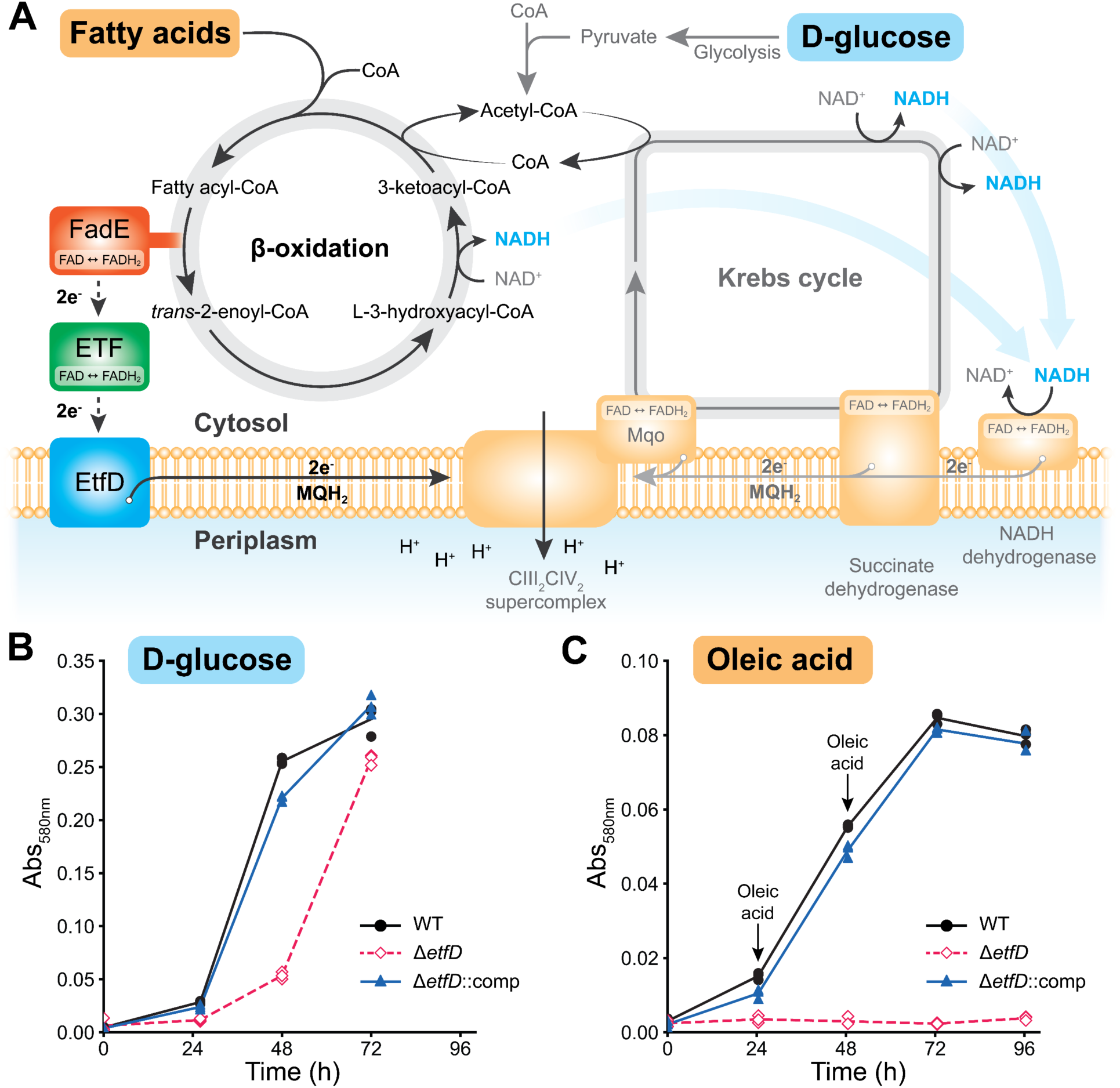
*M. smegmatis* EtfD is essential for growth on fatty acids. A, Schematic representation of the Krebs cycle and β-oxidation in *M. smegmatis*. **B,** Growth of *M. smegmatis* on modified Sauton’s medium supplemented with 11 mM D-glucose. EtfD-3×FLAG (GMC_MSM4) is used as the wild-type (WT) control. **C,** Growth of *M. smegmatis* on modified Sauton’s medium supplemented with 250 µM oleic acid. Each arrow indicates addition of an additional 250 µM oleic acid. Each point shows one of three biological replicates.

The FAD cofactor reduced in β-oxidation is found in a FadE protein (an acyl-CoA dehydrogenase), which transfers its two electrons to a FAD-containing Electron Transfer Flavoprotein (ETF). Mycobacterial ETF (encoded by *rv3028*c and *rv3029c*, also named *etfAB*) and mammalian ETF are homologous. In mammals, an ETF:quinone oxidoreductase (ETF-QO, encoded by the gene *ETFDH*) transfers electrons from ETF to ubiquinone in the mitochondrial inner membrane, which reduces complex III (also known as cytochrome *bc*1) of the ETC. Consequently, mitochondrial ETF and ETF-QO link β-oxidation to the ETC, just as succinate dehydrogenase (Sdh, also known as complex II) (Hägerhäll, 1997) links the Krebs cycle to the ETC. The genes *rv0338c* in *M. tuberculosis* and *MSMEG_0690* in *Mycobacterium smegmatis*, a fast-growing model for mycobacterial energy metabolism, encode a protein named EtfD that has been proposed to serve as an ETF-QO in these organisms (Wischgoll et al., 2005; Agne et al., 2021; Beites et al., 2021). EtfD is expected to transfer electrons from ETF to menaquinone in the plasma membrane, reducing it to menaquinol, which transfers electrons to the complex III2IV2 supercomplex (cyt. *bcc*-*aa*3) or cyt. *bd* (Beites et al., 2021; Zhang et al., 2006). However, mycobacterial EtfD does not appear to be homologous with mammalian ETF-QO.

The *M. tuberculosis* genome encodes many different enzymes that could participate in β- oxidation, including 35 putative acyl-CoA dehydrogenases (Beites et al., 2021; Cole et al., 1998). This diversity leads to an apparent functional redundancy that would make it difficult to target *M. tuberculosis* β-oxidation with inhibitors. However, mycobacterial genomes encode a single ETF and a single EtfD, suggesting a vulnerability in fatty acid catabolism that could be susceptible to inhibition by a single small molecule (Beites et al., 2021). Deletion of EtfD prevents growth of *M. tuberculosis* on fatty acids and cholesterol *in vitro*, and prevents infection in mouse models of TB (Beites et al., 2021). A series of 6,11-dioxobenzo[f]pyrido[1,2-a]indoles (DBPI) compounds have been shown to rely on EtfD to kill *M. tuberculosis* (Székely et al., 2020). The essential nature of EtfD and its lack of homology to human ETF-QO make it an attractive target for treatment of TB. However, the absence of an experimental structure of the protein and the absence of an assay for its activity limit identification and optimization of inhibitors.

In this study, we show that, as in *M. tuberculosis*, deletion of EtfD prevents *M. smegmatis* growth when fatty acids are the sole carbon source available, supporting the use of *M. smegmatis* in the characterization of EtfD inhibitors. We used electron cryomicroscopy (cryo-EM) to determine the *M. smegmatis* EtfD structure, revealing unusual iron-sulfur clusters including a linear [3Fe-4S] cluster that had been produced synthetically and observed spectroscopically but had not been resolved in a protein structure. The iron-sulfur clusters and a *b* heme form a wire that allows electron transfer from ETF in the mycobacterial cytosol to menaquinone in the plasma membrane. Finally, we develop an *in vitro* assay for EtfD activity. The assay confirms the protein’s role in linking fatty acid-oxidation to proton pumping by the ETC and will allow for direct measurement of inhibition of its activity, potentially in high-throughput screening for drug discovery.

## Results

### EtfD is needed for M. smegmatis growth with fatty acids as the sole energy source

Deletion of EtfD in *M. tuberculosis* (*rv0338c*) prevents growth in medium that contains fatty acids or cholesterol (Beites et al., 2021). This observation suggests that a phenotypic screen for growth on medium containing fatty acids may be useful for detecting EtfD inhibitors.

However, *M. tuberculosis* has a doubling time of 24 h and is a human pathogen, complicating its use in high-throughput screens. In contrast, *M. smegmatis* has a doubling time of 3 to 4 h and is nonpathogenic (Reyrat and Kahn, 2001). The value of *M. smegmatis* as a model for energy metabolism in mycobacteria is supported by the phenotypic screen that led to the development of bedaquiline (Andries et al., 2005). Further, *M. tuberculosis rv0338c* and *M. smegmatis MSMEG_0690* are closely related, with 80.2% sequence similarity and 71.2% sequence identity for the folded region of the protein (*MSMEG_0690* residues 1-779, *rv0338c* residues 1-742). For the predicted disordered C-terminal region, the sequence identity and similarity are 36.9% and 39.9%, respectively, with 46.8% gaps. Despite this similarity, whether or not *M. smegmatis* also relies on EtfD for growth on fatty acids is not known. Therefore, we deleted the gene encoding EtfD (*MSMEG_0690*) from *M. smegmatis* using the ORBIT method (Murphy et al., 2018) and tested the ability of the Δ*etfD* strain to grow in liquid medium containing either glucose or oleic acid as the sole carbon source (**Fig. 1B** and **C**, from n=3 biological replicates). With glucose as the sole carbon source, which provides energy through glycolysis and the Krebs cycle, the Δ*etfD* strain demonstrates a slight (∼24 h) growth delay (**Fig. 1B**, *pink*) compared to a wild-type strain (**Fig. 1B**, *black*). A similar delay was reported previously for *M. tuberculosis* Δ*etfD* when grown on glycerol (Beites et al., 2021). The growth defect was rescued by complementing the Δ*etfD* strain with a plasmid that allowed expression of *M. tuberculosis* EtfD (**Fig. 1B**, *blue*). With oleic acid as the sole carbon source, which provides energy through β-oxidation, Δ*etfD M. smegmatis* did not grow (**Fig. 1C**, *pink*), whereas the wild-type strain displayed robust growth (**Fig. 1C**, *black*). Again, this growth defect was rescued by complementation with a plasmid that allowed expression of *M. tuberculosis* EtfD (**Fig. 1C**, *blue*). These data demonstrate that, like *M. tuberculosis*, *M. smegmatis* requires EtfD for growth on fatty acids, and that expression of *M. tuberculosis* EtfD rescues deletion of *M. smegmatis* EtfD. This similarity suggests that *M. smegmatis* may be used in phenotypic screens and characterization of inhibitors of *M. tuberculosis* EtfD.

### Structure of EtfD reveals an electron wire that connects ETF to the menaquinone pool

EtfD has been proposed to transfer electrons from soluble ETF to the membrane-bound menaquinone pool, analogous to human ETF-QO (Beites et al., 2021). However, the two proteins are not homologous and the structural basis for this activity in mycobacteria remains unclear. To investigate this process, we prepared a strain of *M. smegmatis* with a 3×FLAG tag at the C terminus of EtfD, cultured the cells, and isolated membranes. Membranes were then solubilized with the detergent dodecyl maltoside (DDM) and EtfD was purified by affinity chromatography. Cryo-EM of the sample yielded a three-dimensional (3D) map of EtfD at 3.2 Å resolution (**Fig. 2A**, *left*, **Fig. S1**, **Table 1**), with resolution in the soluble region reaching 2.8 Å (**Fig. S2**). This resolution enabled construction of an atomic model for 71% of the 1,042-residue protein (**Fig. 2A**, *right*, **Table 1**). The missing residues in the model correspond to the initial methionine, two loops from residues 348-373 and 621-628, and the ∼30 kDa (residues 778-1042) disordered region at the C terminus of the protein. The ordered portion of EtfD is mostly α-helical (**Fig. 2A**, *right*), consisting of a membrane region comprising five transmembrane α helices and a soluble region that contains two [4Fe-4S] binding domains and two cysteine rich CCG domains (**Fig. 2B**). The two CCG domains coordinate a linear [3Fe-4S] cluster (cluster D1) and a noncubane [4Fe-4S] cluster (cluster D2) (**Fig 2C**, **Movie 1**). The linear [3Fe-4S] cluster D1 is coordinated at each end by Cys503, 538, 584, and 581 (**Fig. 2D**, *top left*). The cluster consists of two perpendicular diamond-shaped planes, matching the structure reported in a chemically synthesized linear [Fe3S4(SPh)4]^3-^ cluster. (Hagen and Holm, 1982). The noncubane [4Fe-4S] D2 cluster is coordinated by Cys637, 672, 673, 709, 712, as well as His 583 (**Fig. 2D**, *top right*). The two [4Fe-4S] domains coordinate two cubane [4Fe-4S] clusters (clusters D3 and D4) (**Fig. 2C**). Cluster D3 is coordinated by Cys311, 412, 415, and 418 while cluster D4 is coordinated by Cys301, 304, 307, and 422 (**Fig. 2D**, *middle*). The membrane embedded region of EtfD contains a heme that, based on the map density and coordinating residues, appears to be a *b* heme (**Fig. 2D**, *bottom left*). His78 and 241 coordinate the metal center of the heme, with Arg137, 150, and 306 forming ionic interactions with the heme’s carboxy groups. Additional density in the map 4- 5 Å from the edge of the heme accommodates what appears to be the napthoquinone head of a menaquinone molecule and likely corresponds to the menaquinone binding site (**Fig. 2D**, *bottom right*). The interaction of this putative menaquinone with the protein is stabilized by aromatic interactions with Phe85 and Phe230. The membrane region of EtfD also appears to interact stably with a detergent-like density that we assigned to a DDM molecule (**Fig. 2A**, *right*).

**Figure 2.**
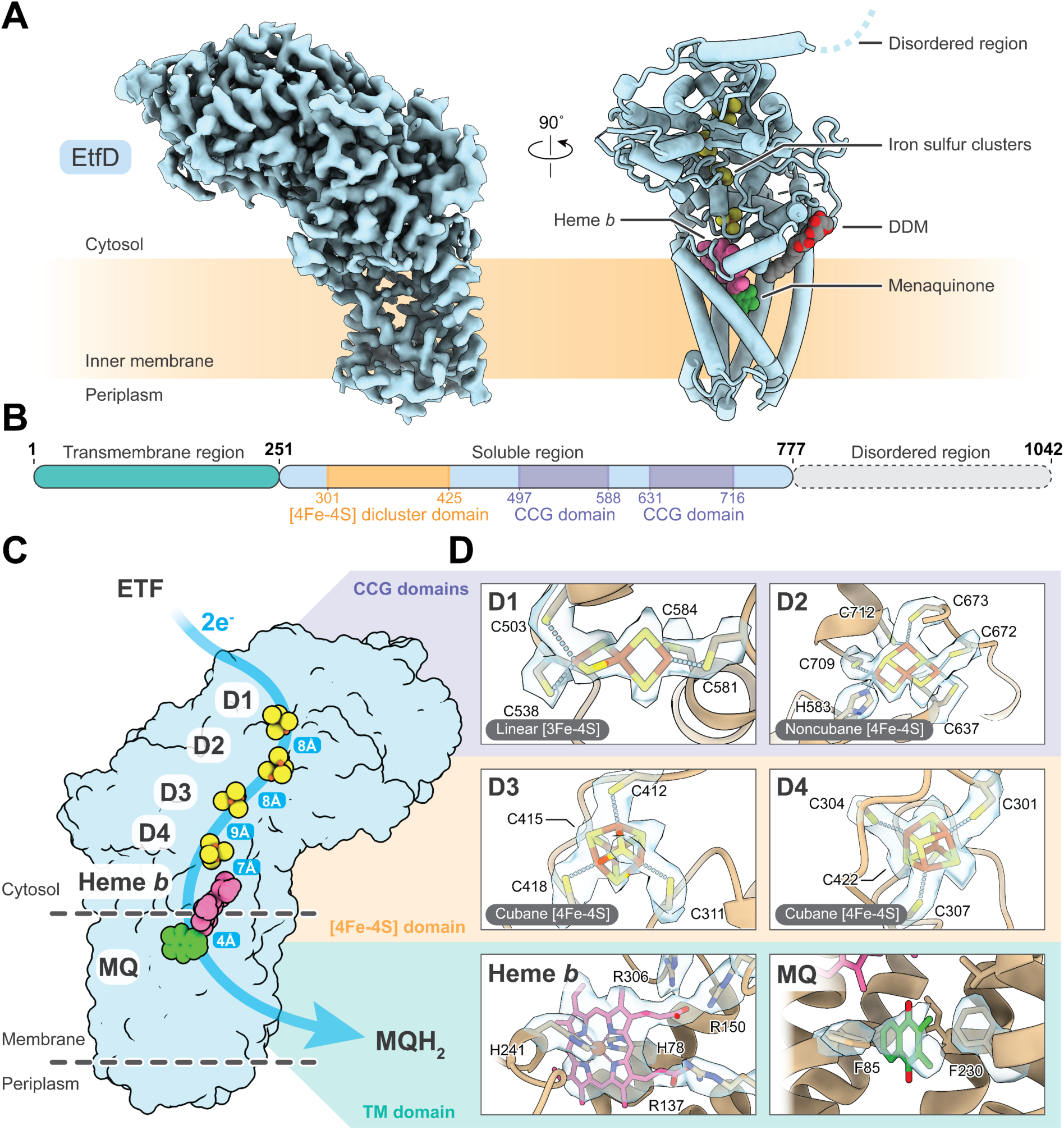
Cryo-EM structure of *M. smegmatis* EtfD. A, Cryo-EM map (*left*) and atomic model (*right*) of EtfD. DDM, dodecyl maltoside. **B,** Schematic representation of EtfD domain organization. Domains were annotated using Pfam (Finn et al., 2008). **C,** Arrangement of redox cofactors in EtfD. A possible electron path from ETF to menaquinone is shown by the blue arrow and the edge-to-edge distance between redox co-factors is indicated. **D,** Coordination of the redox cofactors in EtfD. The cryo-EM map density is shown in blue.

The iron-sulfur clusters, heme, and menaquinone in EtfD are sufficiently close to allow rapid electron transfer between them, with cluster D1 located ∼8 Å away from cluster D2, cluster D2 ∼8 Å from cluster D3, cluster D3 ∼9 Å from cluster D4, cluster D4 ∼7 Å from heme *b*, and heme *b* ∼4-5 Å from the putative menaquinone binding site (**Fig. 2C**). Therefore, the cryo-EM map suggests that EtfD forms a continuous electron wire that can link ETF in the cytosol to the menaquinone pool in the membrane. In this process, electrons could flow sequentially from the FAD in ETF to cluster D1, D2, D3, D4 and the *b* heme, which would reduce menaquinone to menaquinol. However, without knowing where ETF binds on the surface of EtfD, it is not clear which cluster mediates the initial entry of electrons from the FAD of ETF into EtfD.

### Unusual linear and noncubane iron-sulfur clusters in EtfD resemble those in heterodisulfide reductases

The noncubane [4Fe-4S] and linear [3Fe-4S] clusters in EtfD are distinct from iron-sulfur clusters reported in other ETC complexes. A pair of noncubane [4Fe-4S] clusters was observed in the crystal structure of heterodisulfide reductase (Hdr) HdrABC from the anaerobic methanogenic archaeaon *Methanothermococcus thermolithotrophicus* (Wagner et al., 2017; Watanabe et al., 2021). Together, the B and C subunits of the three-subunit HdrABC have the same fold as the soluble region of EtfD (**Fig. 3A***, middle*). The entire EtfD structure also overlays well with an AlphaFold model (Jumper et al., 2021) of the membrane-bound two subunit HdrDE, which is found in archaea such as *Methanosarcina barkeri* (**Fig. 3A**, *right*). The similarity of EtfD and the archaeal Hdrs explains the presence of the noncubane [4Fe-4S] cluster in EtfD (Künkel et al., 1997; Wischgoll et al., 2005). In Hdr, the two noncubane iron-sulfur clusters are solvent-accessible and catalyze the homolytic cleavage of the heterodisulfide formed from coenzyme M and coenzyme B (CoM-S-S-CoB) (**Fig. 3B**, *top).* Each noncubane iron-sulfur cluster harbors a catalytic iron that is coordinated by a bridging cysteine (Cys81 and Cys234) that is displaced in the presence of the ligand (**Fig. 3B**, *bottom, red dashed lines*) (Pelmenschikov et al., 2023). This chemistry allows the noncubane clusters to react with the disulfide bond in the substrate (**Fig. 3B**, *bottom, black dashed lines*) (Wagner et al., 2017), but is also expected to make the clusters in Hdr sensitive to oxygen (Imlay, 2006). In contrast, the iron-sulfur clusters D1 and D2 of EtfD are contained in a cavity that is shielded from the cytosol (**Fig. 3C**, *top*). In cluster D2, the noncubane geometry resembles the noncubane cluster in Hdr but its coordination does not rely on a bridging cysteine. Instead, a histidine at position 583 completes the coordination of the cluster, making it non-catalytic (**Fig. 3C**, *bottom right*). In cluster D1 of EtfD, a threonine at position 539 replaces the cysteine found in Hdr (**Fig. 3C**, *bottom left*).

**Figure 3.**
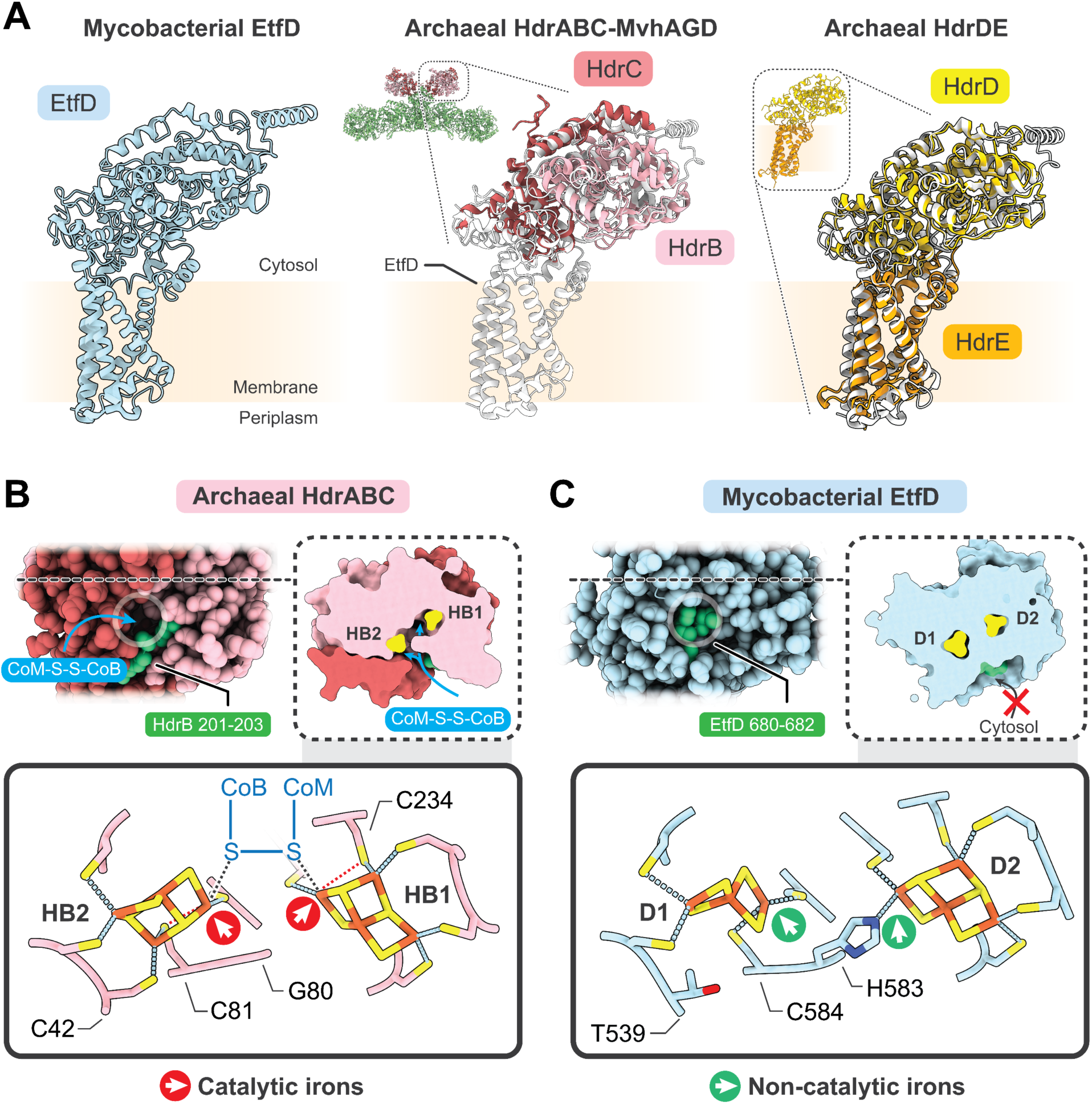
Linear and noncubane iron-sulfur clusters are related to heterodisulfide reductase. A, Comparison of the structure of EtfD (*left*), the archaeal heterodisulfide reductase HdrABC-MvhAGD (*middle,* PDB: 5ODH), and archaeal heterodisulfide reductase HdrDE (*right,* AF-P96796-F1, AF-P96797-F1). **B,** Location (*top*) and coordination (*bottom*) of the catalytic noncubane [4Fe-4S] clusters in HdrABC-MvhAGD. **C,** Location (*top*) and coordination (*bottom*) of the corresponding linear [3Fe-4S] cluster D1 and noncubane [4Fe-4S] cluster D2 in EtfD.

Therefore, EtfD cannot coordinate the fourth iron of a noncubane [4Fe-4S] cluster, and instead uses the cysteine at position 584 to coordinate the third iron and accommodate a linear [3Fe-4S] cluster. (**Fig. 3C**, *bottom left*). The full coordination of the iron-sulfur clusters in EtfD, as well as their shielding from the outside solvent, may indicate adaptation of EtfD to function in aerobic conditions (Imlay, 2006).

### Development of a biochemical assay for mycobacterial EtfD activity

EtfD and its homologues were proposed to link β-oxidation to the ETC by accepting electrons from ETF and using them to reduce the membrane-bound menaquinone pool (Wischgoll et al., 2005; Agne et al., 2021; Beites et al., 2021) (**Fig. 1A**). This role was hypothesized, in part, because of the proximity of the gene encoding EtfD to the genes for the ETF subunits EtfA and EtfB in some bacterial genomes (Wischgoll et al., 2005).

Immunoprecipitation confirmed the interaction of ETF with EtfD, with β-oxidation blocked at the acyl-CoA dehydrogenase step in a Δ*etfD* strain of *M. tuberculosis* (Beites et al., 2021). Based on this model, we developed an assay for EtfD activity (**Fig. 4A**) that resembles previously-developed assays for mycobacterial ATP synthase activity, cyt. *bcc*-*aa*3 activity, and mycobacterial NDH-2 activity (Harden et al., 2024; Liang et al., 2025). In this assay, a fatty acid substrate, acyl-CoA dehydrogenase (FadE) enzyme, ETF, mycobacterial inverted membrane vesicles (IMVs), and the fluorophore 9-amino-6-chloro-2-methoxyacridine (ACMA) are mixed in the wells of a plate in a fluorescence plate reader. The acyl-CoA substrate is oxidized by the FadE enzyme, which transfers electrons to ETF and subsequently EtfD in the IMV membrane. EtfD reduces the membrane-bound menaquinone pool, driving proton pumping by cyt. *bcc*-aa3 and cyt. *bd* and resulting in acidification of the IMV. This acidification causes fluorescence quenching of the ACMA, which recovers when the ΔpH is collapsed by addition of the K^+^/H^+^ antiporter nigericin. Fluorescence quenching and recovery can be detected and quantified using the fluorescence plate reader. We selected butyryl-CoA as the substrate due to its solubility in aqueous solvent, and FadE5 as the acyl-CoA dehydrogenase because its ability to oxidize butyryl-CoA is well characterized (Chen et al., 2020). We prepared *M. smegmatis* strains with 3×FLAG tags at the C termini of the EtfA subunit of ETF and FadE5, and purified endogenous ETF and FadE5 from cultures of these strains (**Fig. S3A**). *M. smegmatis* with no modification of EtfA, EtfB, or EtfD was used to generate wild-type IMVs for assays.

**Figure 4.**
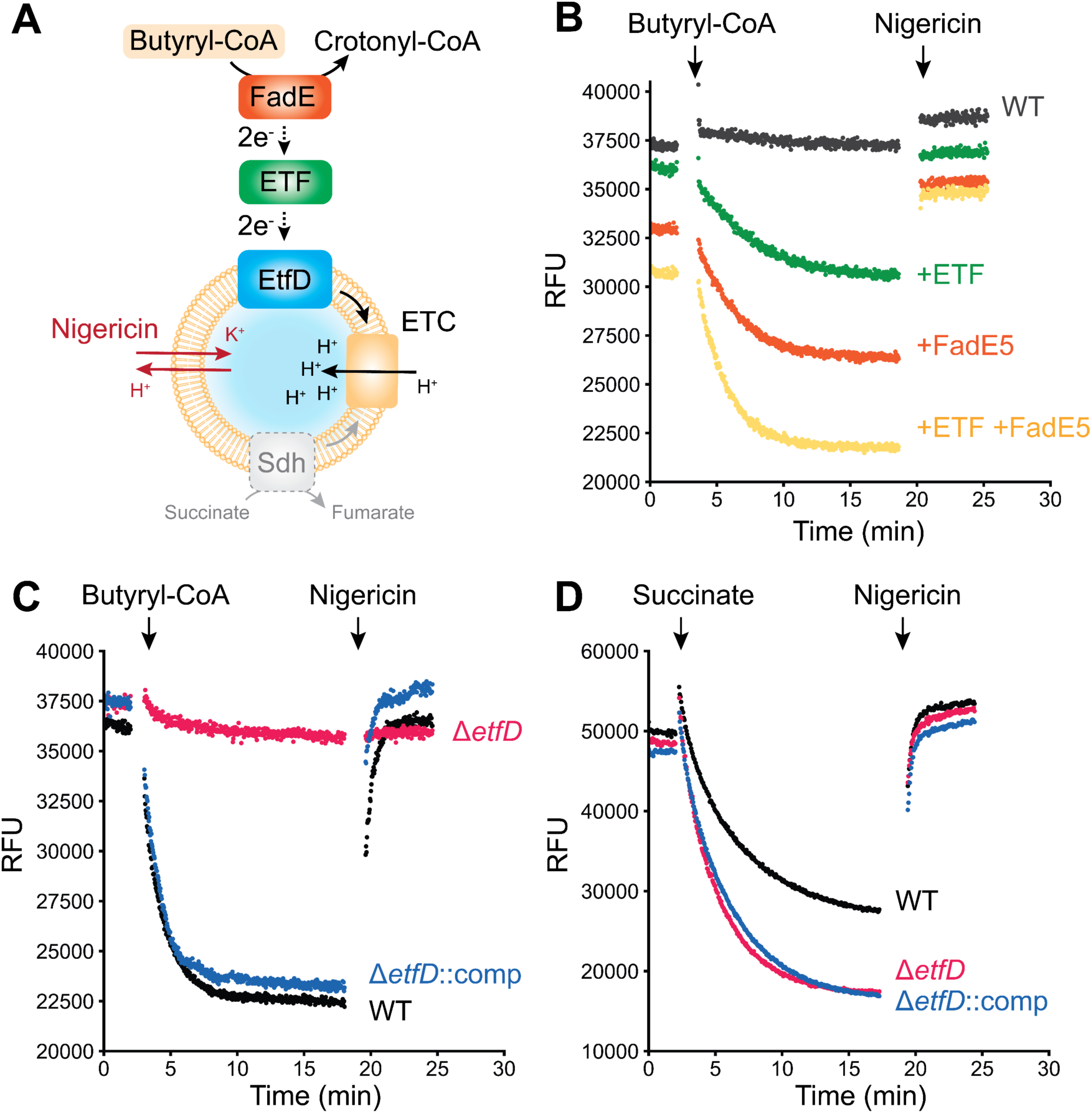
EtfD-mediated proton pumping in the presence of ETF and FadE5. A, Schematic representation of an assay for EtfD activity. **B,** Butyryl-CoA-driven proton pumping activity in IMVs from wild-type *M. smegmatis* (GMC_MSM1), supplemented with either buffer (*black*), ETF (*green*), FadE5 (*orange*), or ETF and FadE5 (*yellow*). **C,** Butyryl-CoA-driven proton pumping activity in the presence of ETF and FadE5 for IMVs prepared from wild-type *M. smegmatis* (*black*), Δ*etfD M. smegmatis* (GMC_MSM9) (*red*), or Δ*etfD M. smegmatis* complemented with *M. tuberculosis* EtfD (GMC_MSM10) (*blue*). **D,** Succinate driven proton pumping activity in IMVs prepared from *M. smegmatis* that is either wild-type (*black*), Δ*etfD* (*red*), or Δ*etfD* complemented with *M. tuberculosis* EtfD (*blue*).

Wild-type *M. smegmatis* IMVs do not display butyryl-CoA dependent acidification (**Fig. 4B**, *grey curve*), likely because little of the soluble proteins FadE and ETF co-purify with these vesicles. However, adding either ETF (**Fig. 4B**, *green*) or FadE5 (**Fig. 4B**, *orange*) leads to butyryl-CoA dependent acidification of IMVs. The observation that adding either ETF or FadE5 allows for activity suggests that trace amounts of each protein contaminate preparations of the other. The extent of acidification was increased by adding both ETF and FadE5 to the assay (**Fig. 4B**, *yellow*). Repeating the experiments that include ETF and FadE5, but using IMVs from a Δ*etfD M. smegmatis* strain shows that acidification is lost completely in the absence of EtfD (**Fig. 4C**, *red*). This acidification is rescued when IMVs are isolated from an *M. smegmatis* strain where the *ΔetfD* mutation is complemented with a plasmid encoding *M. tuberculosis* EtfD (**Fig. 4C**, *blue*). IMVs from wild-type, Δ*etfD*, and Δ*etfD* complemented with a plasmid with *M. tuberculosis* EtfD, all display robust succinate-driven acidification (**Fig. 4D**), where succinate dehydrogenase drives proton pumping by the ETC (Pecsi et al., 2014). These results show that the proton pumping activity measured in the assay is dependent on EtfD, allowing for detection of EtfD activity. Replicates of the assay are shown in Figure S3B-D.

### Effect of DBPI-11626157 on β-oxidation-driven IMV acidification

A series of DBPI compounds was previously shown to rely on EtfD for their bactericidal activity in *M. tuberculosis* (Székely et al., 2020). To investigate the mechanism of action of one of these compounds, DBPI-11626157 (**Fig. S4A**, *left*), we performed assays for butyryl-CoA-driven proton pumping using IMVs from the Δ*etfD* strain complemented with a plasmid for expression of *M. tuberculosis* EtfD. We used these IMVs because DBPIs are active against *M. tuberculosis* but not *M. smegmatis*. DBPI-11626157 at 10 µM (∼6 times its MIC99 of 1.56 µM) inhibited butyryl-CoA-driven proton pumping by 44% (**Fig. S4B** and **D**). To assess whether this inhibition was the result of inhibiting electron transfer from β-oxidation to the ETC, we repeated the experiment but used succinate as the electron donor instead of butyryl-CoA. DBPI-11626157 at 10 µM similarly inhibited succinate-driven proton pumping by 46% (**Fig. S4C** and **D**).

Therefore, DBPI-11626157 does not appear to inhibit EtfD specifically. Interestingly, we noticed that when DBPI-11626157 was present, fluorescence in the assay increased on addition of butyryl-CoA, even when IMV acidification was prevented by including nigericin at the beginning of the experiment (**Fig. S4E**). However, this effect was also observed on addition of butyryl-CoA in the absence of ETF and FadE5, which suggests that it is unrelated to butyryl-CoA mediated-proton pumping (**Fig. S4F**). This increase in fluorescence was not detected when succinate was used as the electron donor (**Fig. S4G**).

## Discussion

*M. tuberculosis* is thought to rely on lipid metabolism during persistence in granulomas (Russell et al., 2009; Wilburn et al., 2018), making EtfD an attractive target for drugs intended to reduce treatment duration. The experiments described here confirm the hypothesis that EtfD forms the electronic link between β-oxidation and the ETC and cryo-EM of EtfD demonstrates that it is structurally unrelated to the mammalian ETF-QO (**Fig. S5A**). We show that *M. smegmatis* provides a model for EtfD activity in mycobacteria, which may enable its use in phenotypic characterization of EtfD inhibitors. Further, we establish a biochemical assay for EtfD activity that can be used in a target-based screen for these compounds.

Disruption of EtfD activity prevents mycobacteria from metabolizing fatty acids and cholesterol as a source of energy, and leads to the accumulation of toxic intermediates in *M. tuberculosis* (Beites et al., 2021). Chemical inhibition of EtfD would not only block β-oxidation but also affect the ability of the ETC to establish a pmf. In *M. tuberculosis*, a Δ*etfD* strain can be complemented with Pox3 *in vitro* (Beites et al., 2021), which has acyl-CoA dehydrogenase activity analogous to FadEs but cannot transfer electrons to the ETC. This observation suggests that the energy contributed to the ETC from β-oxidation is not essential for *M. tuberculosis* survival *in vitro*, but β-oxidation itself, enabled by the oxidation of FadEs by ETF and EtfD, is required (Beites et al., 2021). Whether the contribution of EtfD to the pmf is essential or not *in vivo* remains to be investigated.

While the functional link between ETF and EtfD is clear (Wischgoll et al., 2005; Agne et al., 2021; Beites et al., 2021), ETF was not observed bound to endogenous EtfD in the structure of the protein presented here. This lack of binding could be because ETF shuttles electrons from numerous enzymes to EtfD (Henriques et al., 2021; Beites et al., 2021) and only interacts weakly with EtfD. A weak interaction, which could be lost during protein purification, would explain the lack of EtfD-mediated pumping activity by IMVs that are not supplemented with purified ETF. Alternatively, the structure determined here was obtained under oxidizing conditions, and it is possible that only reduced ETF binds to EtfD. The disordered region of mycobacterial EtfD is not visible in the structure. The length of this region, which includes conserved interspersed repeats (Székely et al., 2020), differs between mycobacterial species (**Fig. S6**). For example, *M. smegmatis* has a ∼30 kDa disordered region, while in *Mycobacterium leprae* it is only ∼13 kDa. An AlphaFold3-predicted structure of the ETF:EtfD complex (**Fig. S5B**) suggests that ETF and EtfD interact through the disordered region of EtfD (**Fig. S5C**).

In the absence of an ETF:EtfD structure, it is not clear which iron-sulfur cluster in EtfD accepts electrons from ETF. The AlphaFold3 model places the FAD moiety of ETF ∼12.5 Å from D1 and ∼11.6 Å from D2 (**Fig. S5D**), which are both sufficiently close for rapid electron transfer. The EtfD homolog FadF in *Bacillus subtilis* is known to be associated with β-oxidation (Matsuoka et al., 2007) and the *Pseudomonas aeruginosa* homolog DgcB participates in dimethylglycine catabolism (Wargo et al., 2008), a function that also involves ETF in mammals (Henriques et al., 2021). However, both FadF and DgcB lack the cysteine residues that coordinate cluster D1 (**Fig. S7**). This observation suggests that FadF and DgcB also accept electrons from ETF and that electrons are transferred from the FAD in ETF directly to the equivalent of cluster D2. Therefore, D2 may also be the initial electron acceptor in EtfD. Interestingly, FadF and DgcB have shorter or entirely absent C-terminal disordered regions (<3 kDa).

The highly unusual linear [3Fe-4S] D1 cluster found in EtfD was first synthesized chemically and later detected spectroscopically in partially unfolded proteins and in the presence of ferrous ammonium sulfate or glutathione (Hagen and Holm, 1982; Kennedy et al., 1984; Gailer et al., 2001; Zhang et al., 2013). Consequently, these structures have been proposed to be degradation products or assembly intermediates of other iron-sulfur clusters (Krebs et al., 2000; Zhang et al., 2013). Linear [3Fe-4S] clusters were recently detected spectroscopically in overexpressed viral proteins under more physiological conditions (Villalta et al., 2023), but it is unknown how these clusters are stabilized in those proteins. The structure presented here demonstrates that a stable linear [3Fe-4S] cluster is found in a natively-folded protein and adopts a geometry similar to the one reported in a synthetic chemical system (Hagen and Holm, 1982). The linear [3Fe-4S] cluster binding motif, which appears related to the noncubane iron-sulfur cluster motif of Hdr, can be found in EtfD homologs from other Actinobacteria (*Streptomyces griseus*) and in diverse phyla such as Acidobacteriota (e.g. *Acidobacterium capsulatum*), Deinococcota (e.g. *Deinococcus proteolyticus*), Spirochaetota (e.g. *Leptospirosa interrogans*), and Gemmatimonadota (e.g. *Gemmatirosa kalamazoonensis*) (**Fig. S7**) (Agne et al., 2021). The properties and role of the linear [3Fe-4S] clusters in these diverse organisms remain to be discovered.

In our assays, DBPI-11626157 shows similar inhibition of butyryl-CoA- and succinate-driven IMV acidification, suggesting that the compound does not inhibit EtfD specifically. DBPI- 11626157 is a menaquinone derivative (**Fig. S4A***, right*), and therefore could interact with other enzymes that use menaquinone. DBPI-11626157 could also uncouple the pmf in IMVs nonspecifically, which could mask a weaker but specific inhibitory activity. The apparent collapse of the pmf in IMVs has been observed for a variety of compounds beyond traditional uncouplers, but is not well understood and does not necessarily translate to disruption of the pmf in live cells (Harrison et al., 2024; Fountain et al., 2025). Therefore, although DBPIs rely on EtfD to kill *M. tuberculosis* (Székely et al., 2020), their mode of action remains to be elucidated. The assay and structure presented here provide a framework for structure-guided drug discovery and development of direct-acting EtfD inhibitors.

## Methods

### Strains

*M. smegmatis* strains GMC_MSM4 (EtfD-3×FLAG), GMC_MSM5 (EtfA-3×FLAG), GMC_MSM9 (Δ*etfD*) and GMC_MSM11 (FadE5-3×FLAG) were generated using the ORBIT method (Murphy et al., 2018). Strains with 3×FLAG tags were prepared using the payload plasmid pSAB41 (Guo et al., 2021) and the deletion strain was prepared using the payload plasmid pKM464 (Addgene #108322) (Murphy et al., 2018). A strain with Δ*etfD* complemented with *M. tuberculosis* EtfD, GMC_MSM10 (Δ*etfD*::*rv0338c*), was generated by transforming GMC_MSM9 with the previously-described integrating plasmid ptb38-*rv0338c* (Beites et al., 2021). GMC_MSM1 (ATP synthase β subunit 3×FLAG) (Guo et al., 2021) was used as a wild-type strain for IMV acidification experiments.

### M. smegmatis growth assay

GMC_MSM4 (EtfD-3×FLAG), GMC_MSM9 (Δ*etfD*), GMC_MSM10 (Δ*etfD*::*rv0338c*) pre-cultures were grown in 7H9 medium supplemented with 0.5% (w/v) fatty acid-free bovine serum albumin (BSA), 0.08% (w/v) NaCl, 0.2% (w/v) D-glucose, 50 µg/mL hygromycin, and 0.05% (v/v) tyloxapol at 37°C with shaking at 180 rpm. Log phase cultures (OD600nm = 0.4 to 0.5) were centrifuged at 6,500 g for 10 min and the bacteria were resuspended in phosphate buffer saline (PBS) to an OD600nm of 0.5. For growth assays, cultures were started at an OD600nm of 0.01 in 2 mL of modified Sauton medium (Beites et al., 2021) supplemented with 0.05% (v/v) tyloxapol, 0.5 % (w/v) BSA, 0.08% (w/v) NaCl, 50 µg/mL hygromycin, and either 0.2% (w/v) D-glucose (∼11 mM) or 250 μM oleic acid as the sole carbon source. Cultures grown with oleic acid were supplemented with an additional 250 μM oleic acid at the ∼24 h and ∼48 h timepoints to allow continued growth while avoiding acute toxicity, as described previously (Beites et al., 2021; Gouzy et al., 2021). Cultures were grown in 2 mL of medium in 14 mL polypropylene round bottom tubes at 37°C with shaking at 180 rpm. Turbidity was assessed by measuring Abs580nm of a 80 µL sample in a transparent flat bottom 96-well microplate (Sarstedt) with a Spectramax M5e plate reader (Molecular Devices).

### Isolation of cytoplasmic and membrane fractions from M. smegmatis

For protein purification and IMV preparation, *M. smegmatis* strains were grown in 25 mL pre-cultures in Middlebrook 7H9 medium supplemented with 0.5% (w/v) BSA, 0.08% (w/v) NaCl, 0.2% (w/v) D-glucose, 50 µg/mL hygromycin, and 0.05% (v/v) Tween 80 at 37°C with shaking at 180 rpm until saturation. From saturated cultures, 2 mL was used to inoculate each of six 1 L cultures grown in Middlebrook 7H9 medium supplemented with 0.08% (w/v) NaCl, 0.2% (w/v) D-glucose, 1% (w/v) tryptone, and 0.05% (v/v) Tween 80. Cells were grown for ∼48 h at 30°C with shaking at 180 rpm. All of the following steps were performed at 4°C or on ice. Cells were collected by centrifugation at 6,500 g for 20 min and resuspended in 150 mL Lysis buffer (50mM Tris-HCl pH 7.5, 150 mM NaCl, 5 mM MgSO4, 5 mM benzamidine hydrochloride, 5 mM aminocaproic acid, 1 mM PMSF). Cells were lysed at 20 kpsi with an Emulsiflex-C3 High-Pressure Homogenizer (Avestin) and then centrifuged at 39,000 g for 30 min. The supernatant was then centrifuged at 200,000 g for 1 h to collect membranes. For GMC_MSM5 (EtfA- 3×FLAG) and GMC_MSM11 (FadE5-3×FLAG) the supernatant was collected to purify ETF and FadE5. For experiments with IMVs and for the purification of EtfD, the membrane fraction was resuspended in 15 mL S-buffer (50 mM Tris-HCl pH 7.5, 150 mM NaCl, 20% (v/v) glycerol, 5 mM MgSO4, 5 mM benzamidine hydrochloride, 5 mM aminocaproic acid, 1 mM PMSF), flash frozen in liquid nitrogen, and stored at -80°C. For IMV assays, resuspended membranes were aliquoted before freezing.

### IMV acidification assay

Proton pumping was assayed using IMVs harvested from GMC_MSM1, GMC_MSM9, and GMC_MSM10, with GMC_MSM1 acting as the wild-type control. Cells were grown for either 40 or 48 h. The assay was performed in IMV ACMA buffer (10 mM HEPES-KOH pH 7.5, 100 mM KCl, 5 mM MgCl2, 3 µM ACMA) at a final volume of 160 µL. IMVs were used at a final concentration of ∼1-1.5 mg/mL when butyryl-CoA was the electron donor, and diluted 16-fold when succinate was the electron donor. When comparing strains IMVs were diluted to normalize protein concentration. When used, ETF and FadE5 were added to IMVs at a final concentration of 2 µM each, or substituted with FLAG wash buffer 2 (50 mM Tris-HCl pH 7.4, 150 mM NaCl, 20% (v/v) glycerol, 5 mM MgSO4, 5 mM benzamidine hydrochloride, 5 mM aminocaproic acid). In inhibition assays, DBPI-11626157 was first dissolved in DMSO at a concentration of 10 mM, stored at -20°C and sonicated before use in a bath sonicator. The compound was added to the assay at a final DMSO concentration of 2%. The reaction was started with 2 mM (final concentration) butyryl-CoA lithium salt hydrate (MilliporeSigma) in MilliQ water (pH ∼ 4/5), or 5 mM disodium succinate (pH ∼8). To dissipate the proton gradient, nigericin (3.28 µL of a 50 µM stock in 1% ethanol) was added to the 160 µL reaction 15 min after the addition of the electron donor, reaching a final concentration of 1 µM. Black round bottom 96-well microplates (BRANDTECH scientific) were used, and the fluorescence was excited at 410 nm and monitored at 480 nm with a BioTek Synergy Neo2 Multi-mode Assay Microplate reader (Agilent technologies). The difference in relative fluorescence units before and after addition of nigericin was calculated to quantify proton pumping activity. All values were divided by the mean fluorescence recovery of the DMSO controls to obtain relative activity. Importantly, identical assays performed with butyryl-CoA lithium salt hydrate from a different vendor (CHEMIMPEX) had a pH of ∼7 when dissolved with MilliQ water and showed acidification of wild-type IMVs and Δ*etfD* IMVs both with and without ETF and FadE5, neither of which are expected to be capable of butyryl-CoA-driven acidification. This unexpected activity may be due to the presence of contaminants in the substrate that can donate electrons to other complexes of the ETC. Consequently, great care must be taken in identifying a suitable source of butyryl-CoA for the assay.

### Protein purification

To purify EtfD, membranes were solubilized in 1% (w/v) dodecyl maltoside (DDM) for 1 h. Insoluble material was removed by centrifugation at 200,000 g for 1h. The supernatant was collected and filtered through a 0.45 µm filter before being applied to a column of 2 mL M2 affinity matrix (Sigma) previously equilibrated with FLAG wash buffer 1 (50 mM Tris-HCl pH 7.4, 150 mM NaCl, 15% (v/v) glycerol, 0.05% (w/v) DDM, 5 mM benzamidine hydrochloride, 5 mM aminocaproic acid). The column was then washed with ten column volumes of FLAG wash buffer 1, before protein was eluted with three column volumes of FLAG wash buffer 1 containing 150 μg/mL 3×FLAG peptide. The sample was then concentrated with a 30 kDa cutoff concentrator (Sigma) to 500 µL before being loaded onto a Superose 6 Increase 10/300 gel filtration column (GE Healthcare) equilibrated in gel filtration buffer (50 mM Tris-HCl pH 7.4, 150 mM NaCl, 15% glycerol, 0.05% (w/v) DDM). Fractions containing EtfD were pooled and concentrated to ∼12 μL with a 100 kDa cutoff concentrator (Sigma) for cryo-EM sample preparation.

To purify ETF and FadE5, the cytoplasmic fraction obtained after removing membranes from GMC_MSM5 and GMC_MSM11 cells was filtered with a 0.45 µM filter and applied to a column of 2 mL of M2 affinity matrix previously equilibrated using FLAG wash buffer 2 (50 mM Tris-HCl pH 7.4, 150 mM NaCl, 20% (v/v) glycerol, 5 mM MgSO4, 5 mM benzamidine hydrochloride, 5 mM aminocaproic acid). The column was then washed with ten column volumes of FLAG wash buffer 2, before bound protein was eluted with three column volumes of FLAG wash buffer 2 containing 150 μg/mL 3×FLAG peptide. The eluted protein was then concentrated to ∼30-60 µM with a 30 kDa cutoff concentrator.

### Cryo-EM sample preparation

Holey gold grids were made as described previously (Marr et al., 2014). Prior to grid freezing, glycerol was removed from the sample with a Zeba spin desalting column (Thermo Fischer) equilibrated in freeze buffer (50 mM Tris-HCl pH 7.4, 150 mM NaCl, 0.05% (w/v) DDM). Grids were glow discharged in air for 2 min and 2 µL of sample was applied to each grid in EM GP2 plunge freezer (Leica) at 4°C and 90% humidity. Grids were blotted for 1 s before freezing in liquid ethane.

### Data collection

Cryo-EM data was acquired with a 300kV Titan Krios G3 electron microscope equipped with a Falcon 4i camera (Thermo Fischer Scientific). Data collection was automated with the EPU software package. 11,952 movies were collected in EER format (Guo et al., 2020) at 120,000× magnification, corresponding to calibrated pixel size of 0.64 Å. The exposure rate of 7.7 e^-^/pixel/s with a total exposure of ∼70 e^-^/Å^2^.

### Image analysis

cryoSPARC v.4 (Punjani et al., 2017) was used for image analysis. Movies were first aligned using patch motion correction and contrast transfer function (CTF) parameters were estimated in patches. Blob picking and 2D classification were used to generate 2D templates for template-based particle selection. Template selection yielded 1,588,703 particle images, which were extracted using a box size of 512×512 pixels, and Fourier cropped to 208×208 pixels. The dataset was curated using 2D classification and several rounds of ab-initio and heterogenous refinement, yielding 152,064 particle images. Particle images were then re-extracted using a box size of 512×512 pixels and Fourier cropped to 320×320 pixels. Non-uniform refinement (Punjani et al., 2020) and CTF refinement yielded a map of EtfD at 3.1 Å resolution. This resolution was improved to 2.9 Å with two rounds of reference-based motion correction and CTF refinement.

Local refinement of the soluble region of EtfD yielded a map at 2.8 Å resolution. 3D classification was used to separate EtfD with an intact membrane region from EtfD particles lacking the heme density. Non-uniform refinement of 48,099 intact particles from the 3D class that included the heme density yielded a 3.1 Å resolution map for the intact protein. However, non-uniform refinement of 48,053 particles images re-extracted from pre-processed movies resulted in a map that was at 3.2 Å resolution, but which showed better-defined map densities for the membrane-embedded region of EtfD.

### Atomic model building and refinement

An AlphaFold 3 (Jumper et al., 2021) prediction of the *M. smegmatis* EtfD structure (AFDB ID number: AF-A0QQB0-F1-v4) was fitted as a rigid body in the experimental density map for EtfD using UCSF Chimera (Goddard et al., 2007). The model was adjusted with Coot (Emsley and Cowtan, 2004) and refined with ISOLDE (Croll, 2018) and PHENIX (Afonine et al., 2018). Model statistics were evaluated with EMRinger (Barad et al., 2015) and Molprobity (Chen et al., 2010). Amino acid residues where the map lacked density for the side chain were truncated at the β carbon in the final model. Figures and movies were made using UCSF ChimeraX (Goddard et al., 2018) and Microsoft Clipchamp. Restraint files for the noncubane [4Fe-4S] and linear [3Fe- 4S] clusters were generated with ELBOW (Moriarty et al., 2009) using the computationally-predicted structure of an open noncubane cluster (Pelmenschikov et al., 2023) and the crystal structure of the synthetic linear [Fe3S4(SPh)4]^3-^ cluster (Hagen and Holm, 1982).

### Sequence alignments

Sequences were aligned using Clustal Omega (Goujon et al., 2010; Sievers et al., 2011) and analyzed with Jalview (Waterhouse et al., 2009).

## Data availability

Cryo-EM maps and the final atomic model are available through the Electron Microscopy Databank with accession codes EMD-70545 and EMD-70546, and Protein Databank with accession code 9OJN.

## Acknowledgements

We thank Yingke Liang for advice on protein purification strategies. GMC was supported by a Mary H. Beatty Fellowship. JLR was supported by the Canada Research Chairs program. Research was funded by the Canadian Institutes of Health Research Project (grant PJT191893). SE and DS were supported by the National Institutes of Health (grant AI143575) and the Gates Foundation (grants INV-055896 and INV-055894). Cryo-EM data were collected at the Toronto High-Resolution High Throughput cryo-EM facility, supported by the Canada Foundation for Innovation and Ontario Research Fund. Assays were performed using infrastructure from the Structural and Biophysical Core Facility at The Hospital for Sick Children,

## Statement of contributions

GMC conceived the project, designed and performed assays, purified protein, and collected and analyzed cryo-EM data. VM and SC provided the compound DBPI-11626157 and advised on its use. SE and DS provided guidance on growth of *Mycobacterium smegmatis* with fatty acids as the energy source and provided the plasmid for expression of *M. tuberculosis* EtfD. JLR supervised the research, provided advice on the design of experiments, and coordinated the project. GMC and JLR wrote the manuscript and prepared the figures with input from the other authors.

## Conflict of interests

The authors declare that they have no conflict of interest.

---

**Figure S1.**
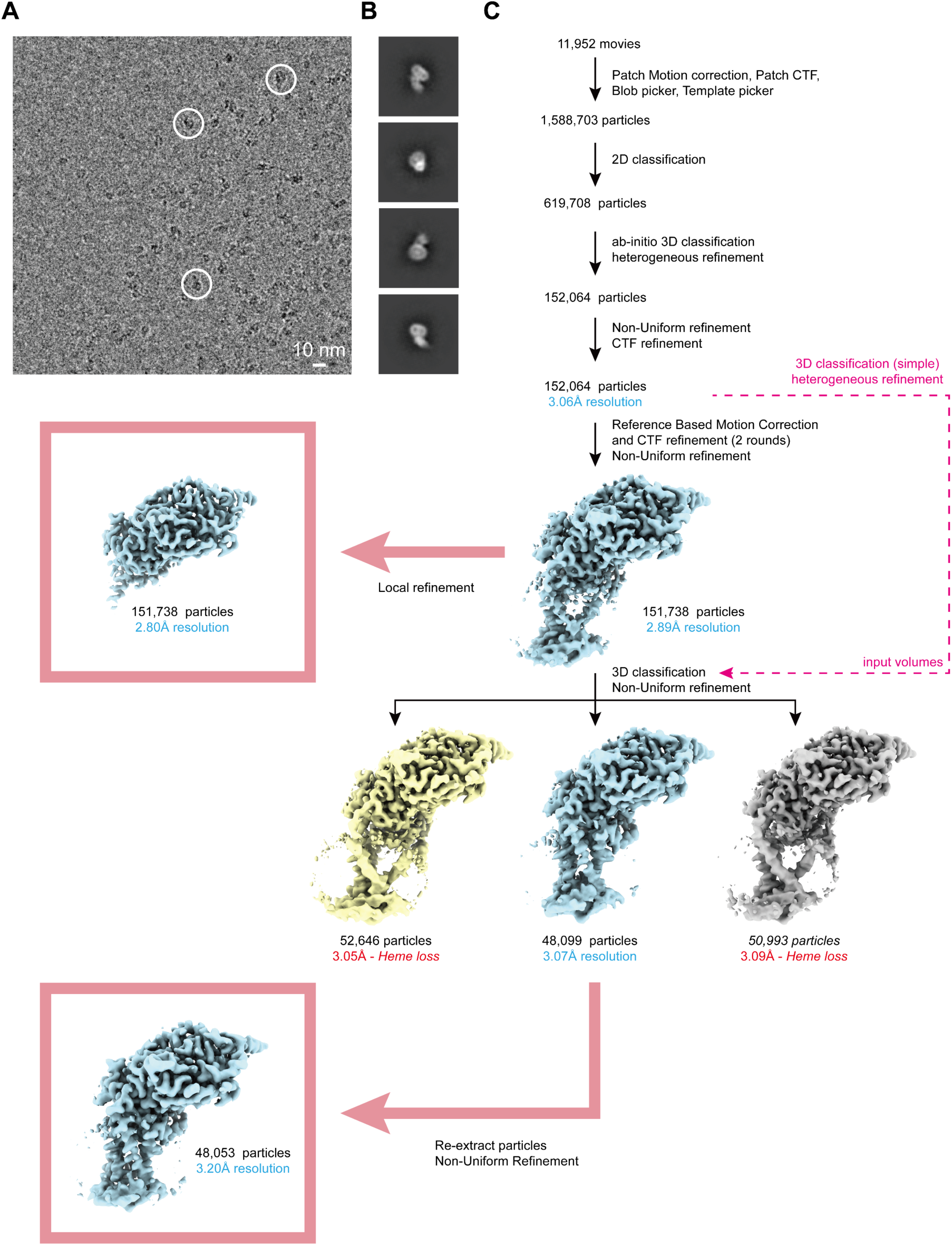
Cryo-EM workflow. A, Representative micrograph. Example particle images are circled. **B,** 2D class averages of EtfD. **C,** Simplified cryo-EM workflow.

**Figure S2.**
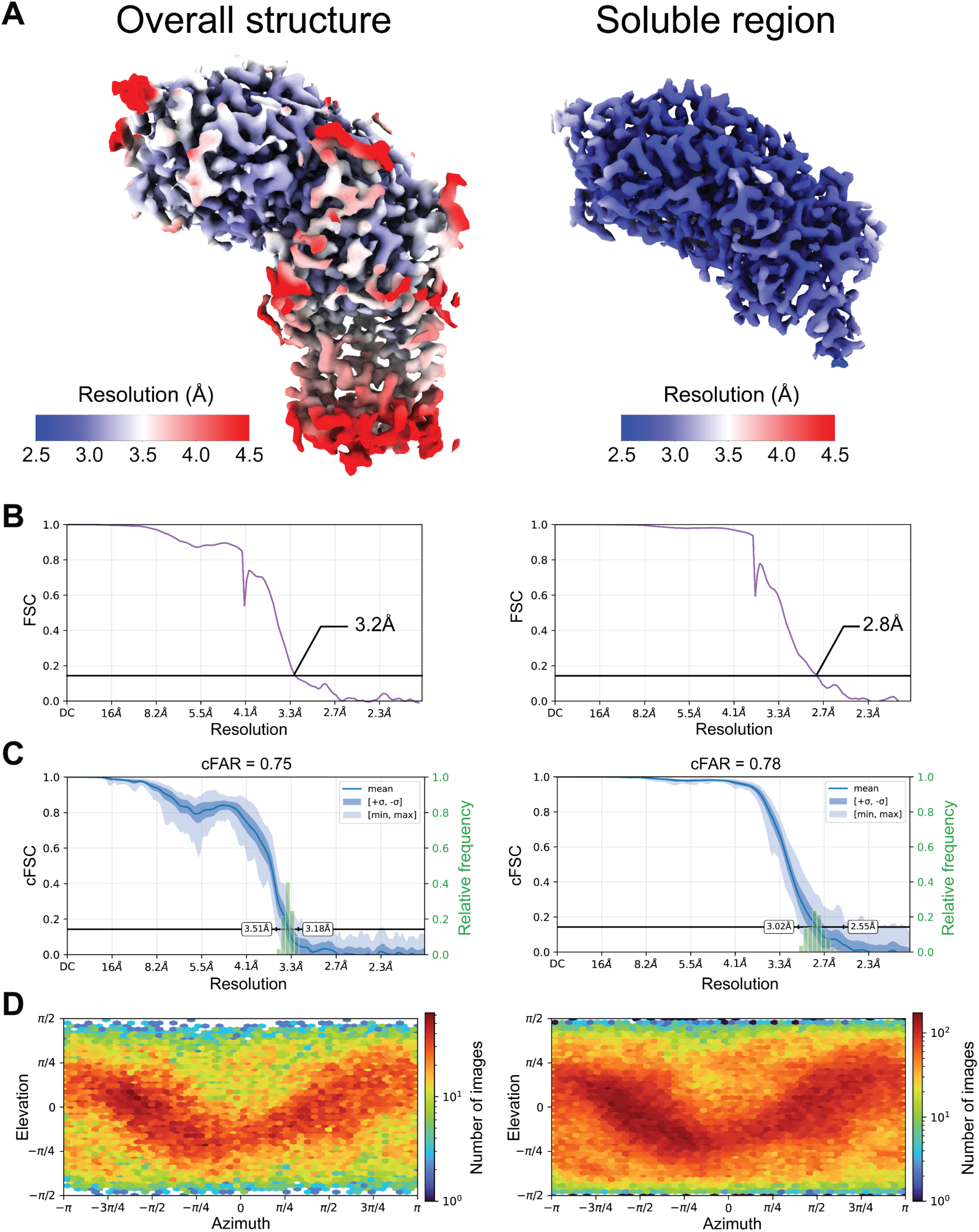
Cryo-EM map validation. A, Local resolution map for the full EtfD structure (*left*) and the locally refined soluble region of EtfD (*right*). **B,** Fourier Shell Correlation (FSC) curve obtained after gold-standard refinement and corrected for the effects of masking. **C,** Conical FSC (cFSC) plots and their associated cFSC area ratio (cFAR) scores. **D,** Particle orientation distribution plots.

**Figure S3.**
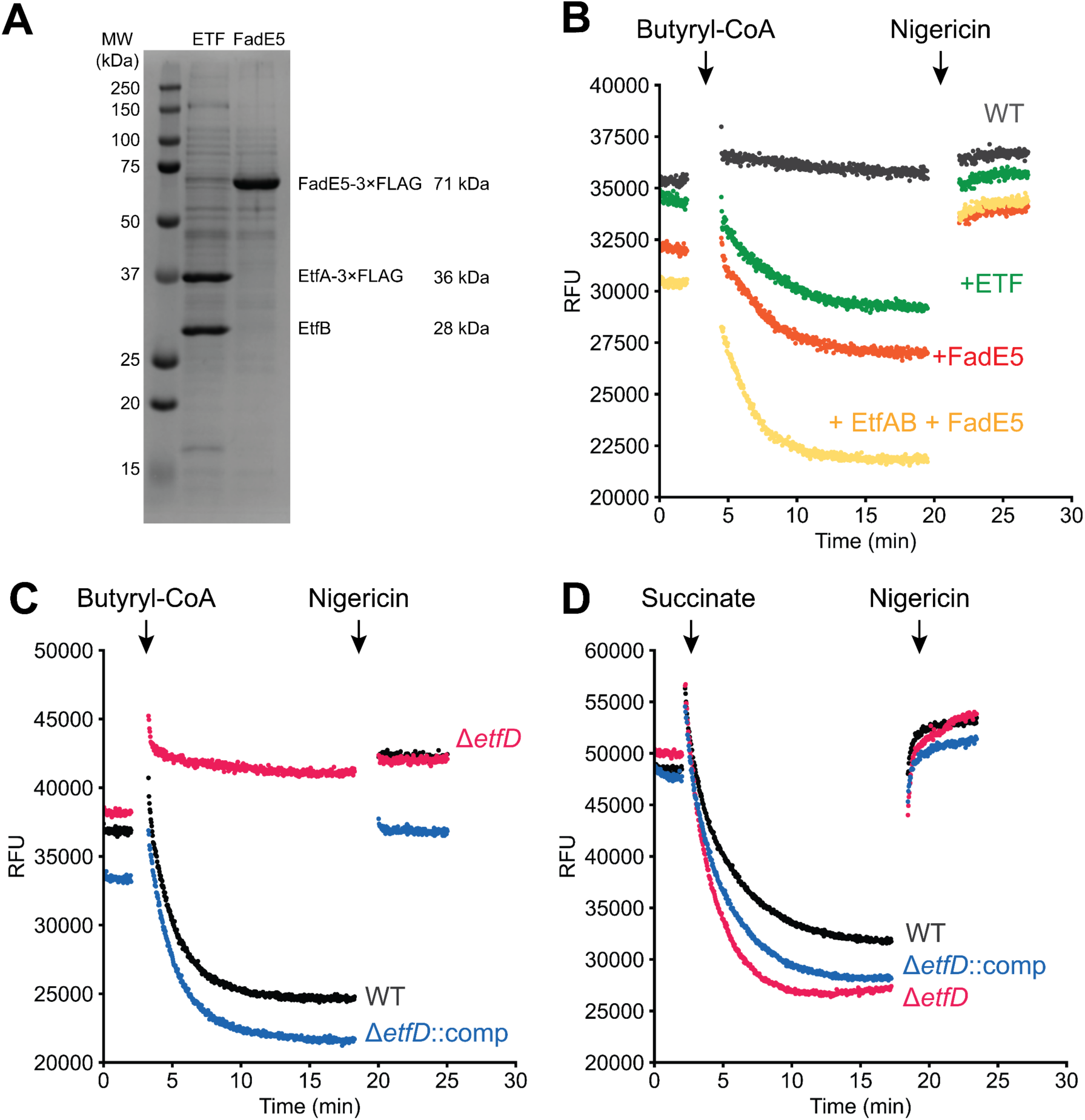
Assay replicates. A, SDS-PAGE gels of purified ETF and FadE5. **B,** Butyryl-CoA- driven proton pumping activity in IMVs prepared from wild-type *M. smegmatis* (GMC_MSM1) supplemented with either buffer (*black*), ETF (*green*), FadE5 (*orange*), or ETF and FadE5 (*yellow*). **C,** Butyryl-CoA-driven proton pumping activity in the presence of ETF and FadE5 for IMVs prepared from *M. smegmatis* that is either wild-type (*black*), Δ*etfD* (*red*) (GMC_MSM9), or Δ*etfD* complemented with *M. tuberculosis* EtfD (GMC_MSM10). **D,** Succinate-driven proton pumping activity in IMVs prepared from *M. smegmatis* that is either wild-type (*black*), Δ*etfD* (*red*), or Δ*etfD* complemented with *M. tuberculosis* EtfD (*blue*).

**Figure S4.**
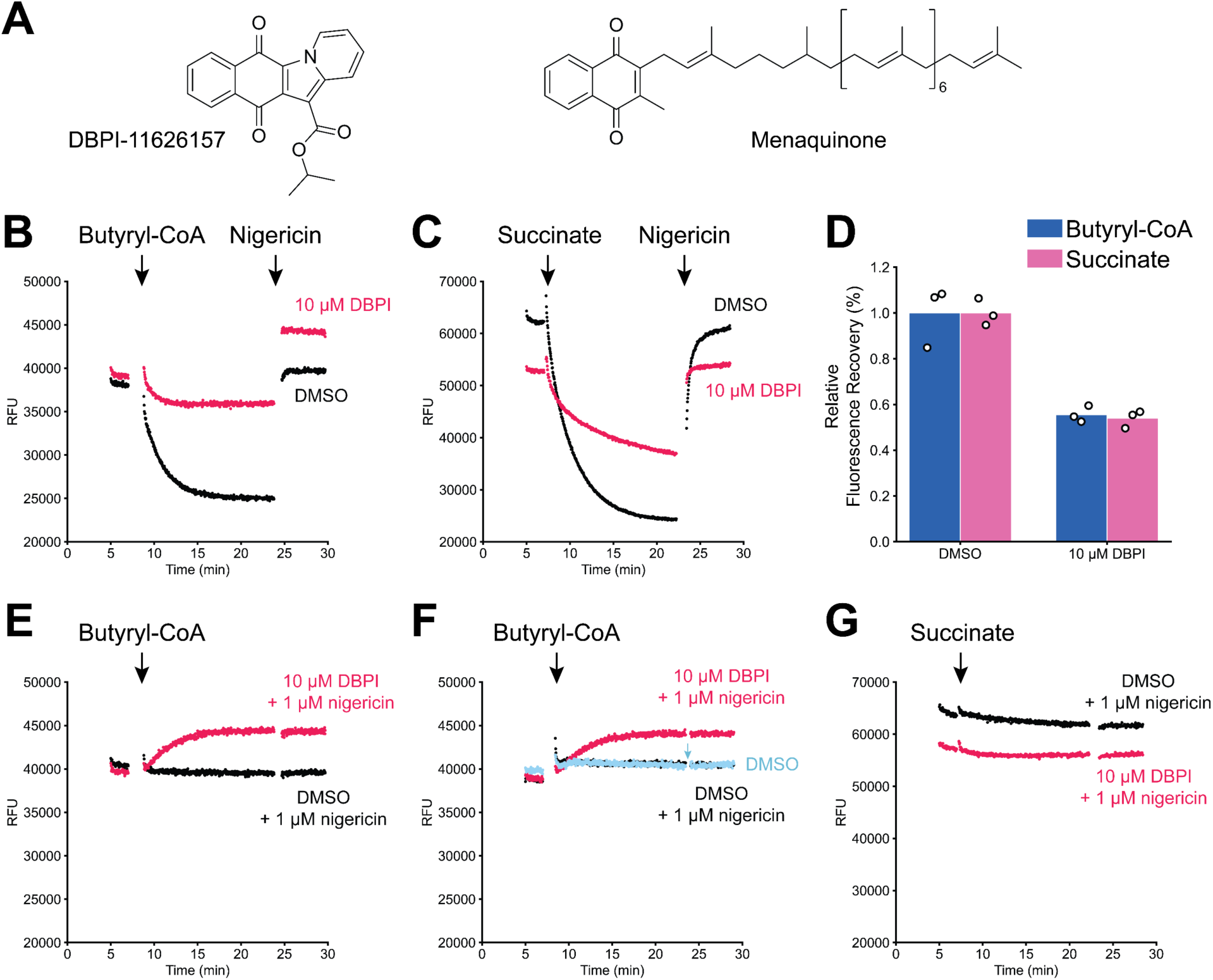
Effect of DBPI-11626157 on butyryl-CoA-driven and succinate-driven IMV acidification. A, Chemical structure of DBPI-11626157 (*left*) and menaquinone (*right*). **B,** Butyryl-CoA-driven acidification of Δ*etfD*::*rv0338c* IMVs in the presence of 2 μM ETF and 1 µM FadE5, with 10 μM DBPI-11626157 (*red*) or DMSO (*black*). Data is representative of three replicates. **C,** Succinate-driven acidification of Δ*etfD*::*rv0338c* IMVs with 10 μM DBPI- 11626157 (*red*) or DMSO (*black*). Data is representative of three replicates. **D,** Quantification of butyryl-CoA-driven and succinate-driven IMV acidification assays. **E,** Addition of butyryl-CoA to Δ*etfD*::*rv0338c* IMVs in the presence of 1 μM nigericin, 2 μM ETF, and 1 µM FadE5, with 10 μM DBPI-11626157 (*red*) or DMSO (*black*). Data is representative of three replicates. **F,** Addition of butyryl-CoA to Δ*etfD*::*rv0338c* IMVs without FadE5 or ETF with 10 μM DBPI- 11626157 and 1 μM nigericin (*red*), DMSO and 1 μM nigericin (*black*), or DMSO alone (*blue*). Nigericin (1 μM) was added to the DMSO-only well during the assay (*blue arrow*). Data is representative of two replicates. **G,** Addition of succinate to Δ*etfD*::*rv0338c* IMVs in the presence of 1 μM nigericin with 10 μM DBPI-11626157 (*red*) or 1 μM nigericin and DMSO (*black*). Data is representative of three replicates.

**Figure S5.**
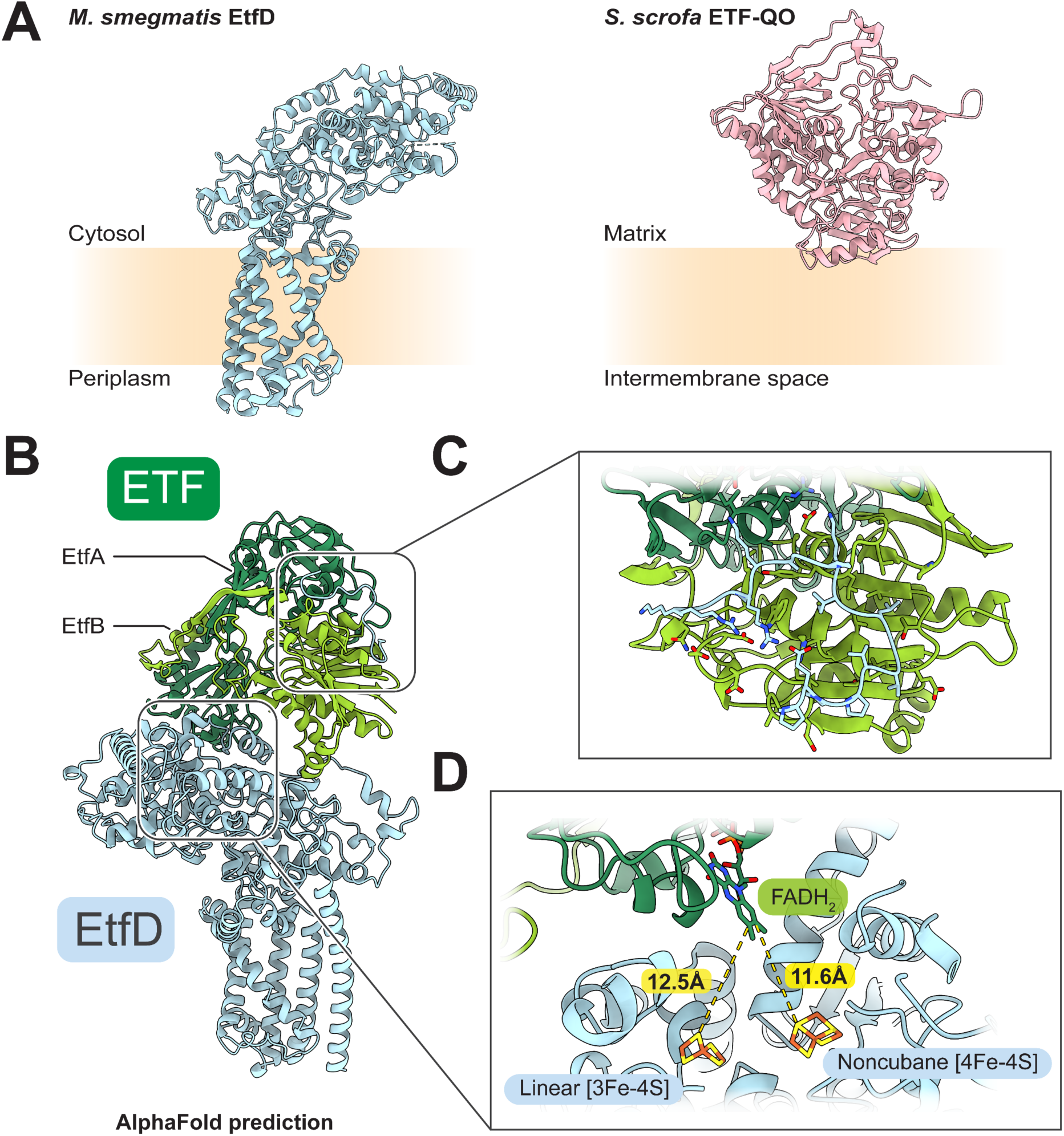
AlphaFold prediction of the ETF:EtfD complex. A, Comparison of EtfD from *M. smegmatis* with ETF-QO oxidoreductase from *Sus scrofa* (PDB: 2GMH) **B,** Overview of the interaction between ETF (*green*) and EtfD (*blue*) predicted by AlphaFold 3. The ∼30 kDa disordered region of EtfD is not shown for clarity. **C,** Close-up view of the interaction predicted by AlphaFold 3 between the disordered region of EtfD and ETF. **D,** Location of the docked FAD cofactor of ETF from PDB 1EFV, relative to cluster D1 and D2 of EtfD.

**Figure S6.**
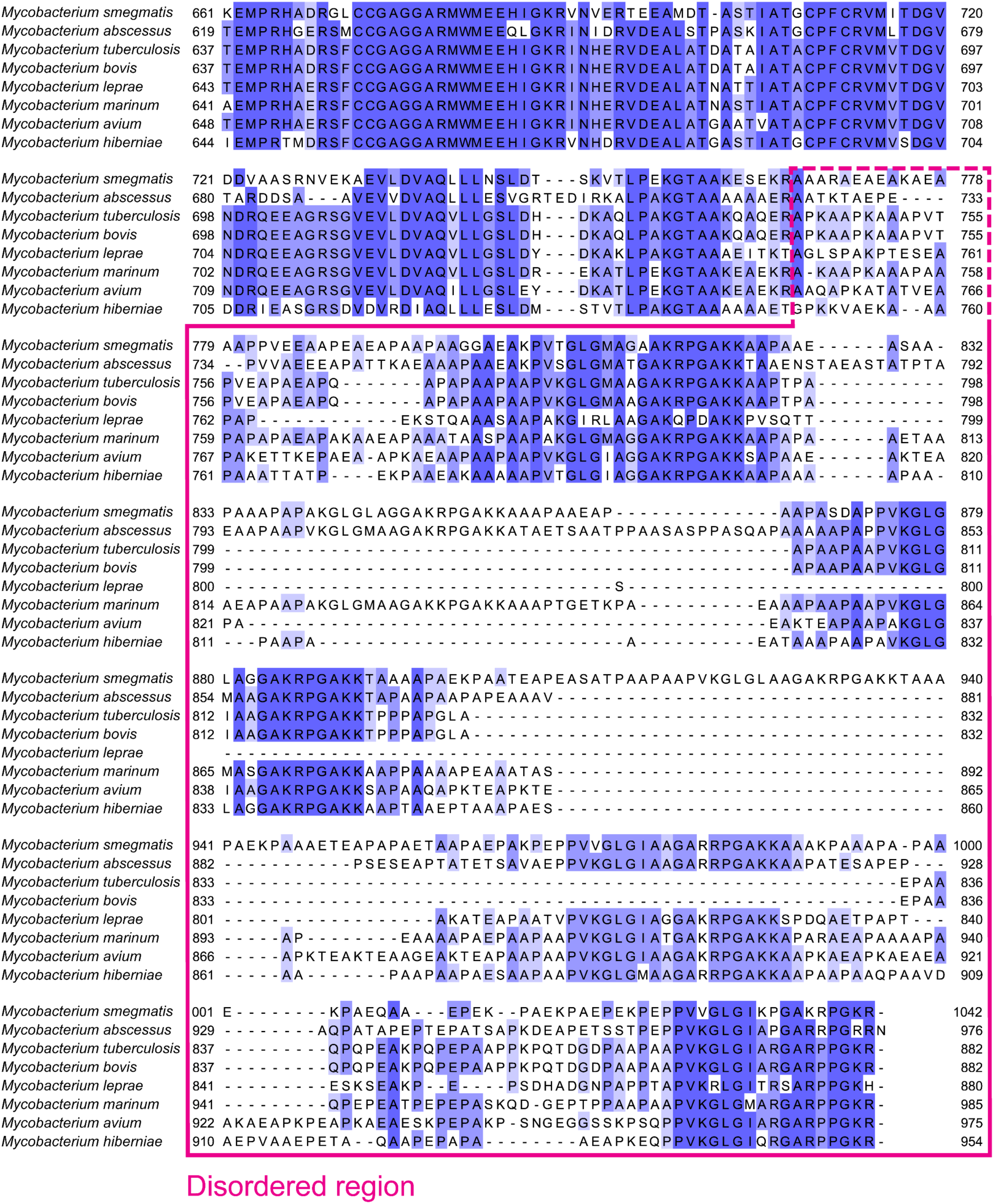
Sequence alignment of the disordered region of mycobacterial EtfD homologs. Sequences are colored by conservation. The disordered region is indicated by a pink box. The start of the disordered region differs between species (dashed line).

**Figure S7.**
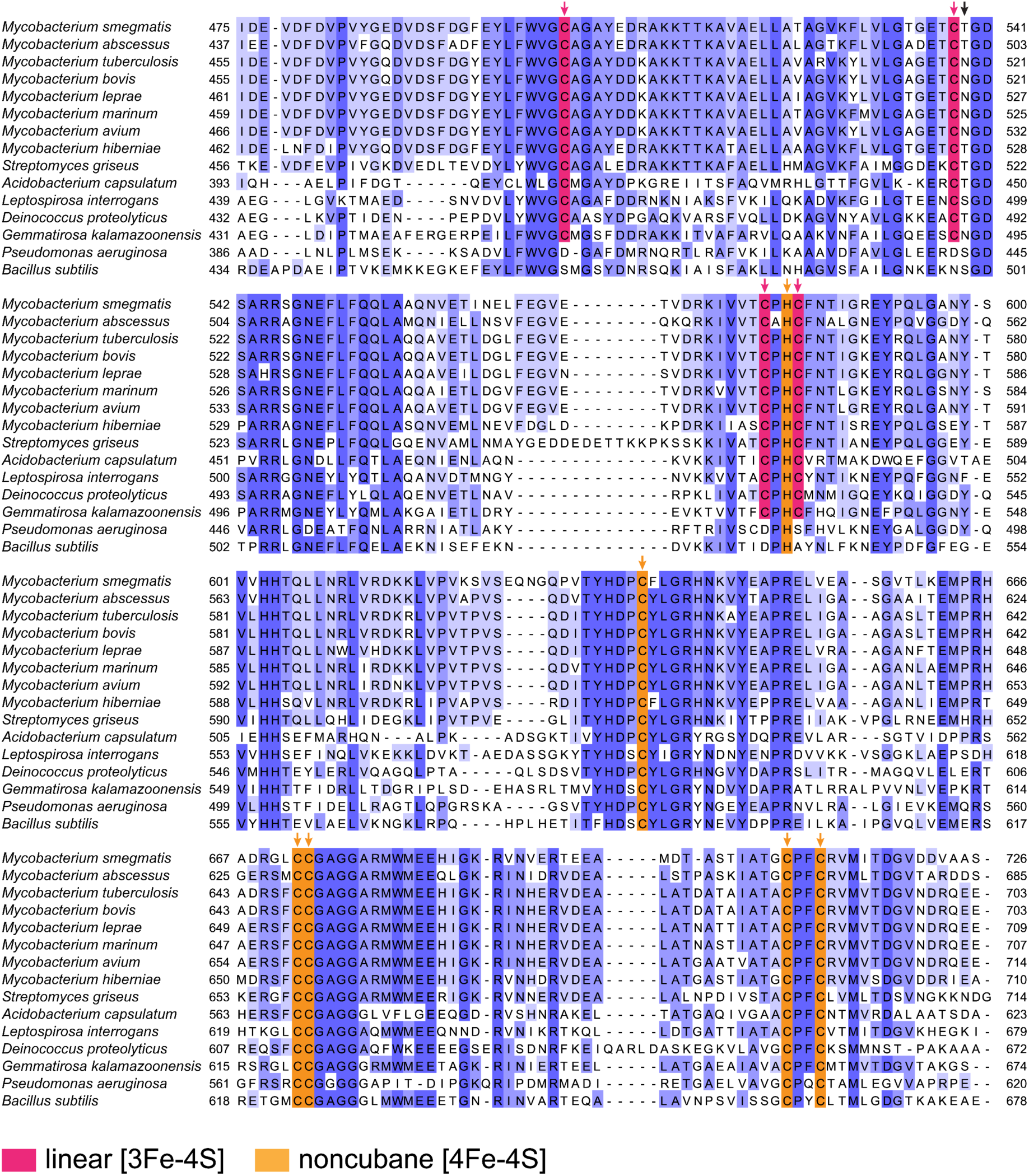
Sequence alignment of the CCG domains of selected EtfD homologs. Coordinating residues are shown with red (linear [3Fe-4S] cluster) and orange (noncubane [4Fe- 4S] cluster) arrows. The black arrow shows the position of the supernumerary cysteine in Hdr. Sequences are colored by conservation.

**Supplementary Table 1.**
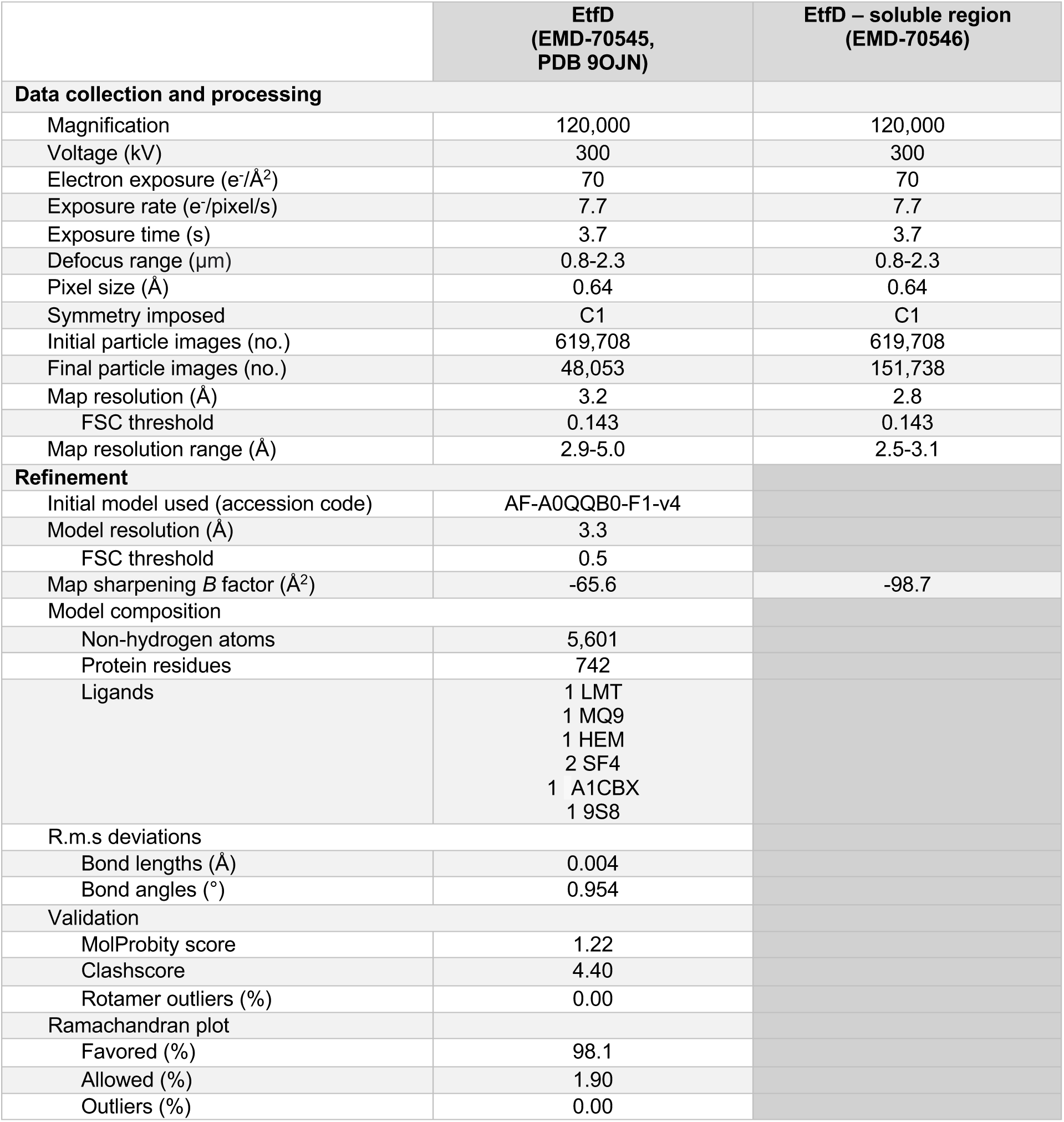
Cryo-EM data collection, refinement, and validation statistics.

|  | **EtFD**<br>**(EMD-70545,**<br>**PDB 9OJN)** | **EtFD – soluble region**<br>**(EMD-70546)** |
| --- | --- | --- |
| **Data collection and processing** |  |  |
| Magnification | 120,000 | 120,000 |
| Voltage (kV) | 300 | 300 |
| Electron exposure (e<sup>-</sup>/Å<sup>2</sup>) | 70 | 70 |
| Exposure rate (e<sup>-</sup>/pixel/s) | 7.7 | 7.7 |
| Exposure time (s) | 3.7 | 3.7 |
| Defocus range (μm) | 0.8-2.3 | 0.8-2.3 |
| Pixel size (Å) | 0.64 | 0.64 |
| Symmetry imposed | C1 | C1 |
| Initial particle images (no.) | 619,708 | 619,708 |
| Final particle images (no.) | 48,053 | 151,738 |
| Map resolution (Å) | 3.2 | 2.8 |
| FSC threshold | 0.143 | 0.143 |
| Map resolution range (Å) | 2.9-5.0 | 2.5-3.1 |
| **Refinement** |  |  |
| Initial model used (accession code) | AF-A0QQB0-F1-v4 |  |
| Model resolution (Å) | 3.3 |  |
| FSC threshold | 0.5 |  |
| Map sharpening *B* factor (Å<sup>2</sup>) | -65.6 | -98.7 |
| **Model composition** |  |  |
| Non-hydrogen atoms | 5,601 |  |
| Protein residues | 742 |  |
| Ligands | 1 LMT<br>1 MQ9<br>1 HEM<br>2 SF4<br>1 A1CBX<br>1 9S8 |  |
| **R.m.s deviations** |  |  |
| Bond lengths (Å) | 0.004 |  |
| Bond angles (°) | 0.954 |  |
| **Validation** |  |  |
| MolProbity score | 1.22 |  |
| Clashscore | 4.40 |  |
| Rotamer outliers (%) | 0.00 |  |
| **Ramachandran plot** |  |  |
| Favored (%) | 98.1 |  |
| Allowed (%) | 1.90 |  |
| Outliers (%) | 0.00 |  |

## Notes

### Competing Interest Statement

The authors have declared no competing interest.

